# IPSILESIONAL HIPPOCAMPAL GABA IS ELEVATED AND CORRELATES WITH COGNITIVE IMPAIRMENT AND MALADAPTIVE NEUROGENESIS AFTER CORTICAL STROKE IN MICE

**DOI:** 10.1101/2022.11.02.514653

**Authors:** C Torres-López, MI Cuartero, A García-Culebras, J de la Parra, ME Fernández-Valle, M Benito, S Vázquez-Reyes, T Jareño-Flores, O Hurtado, MS Buckwalter, JM García-Segura, I Lizasoain, MA Moro

## Abstract

**Background:** Cognitive dysfunction is a frequent stroke sequela but its pathogenesis and treatment remain unresolved. Involvement of aberrant hippocampal neurogenesis and maladaptive circuitry remodelling has been proposed but their mechanisms are unknown. Our aim was to evaluate potential underlying molecular/cellular events implicated.

**Methods:** Stroke was induced by permanent occlusion of the middle cerebral artery (MCAO) in 2-month-old C57BL/6 male mice. Hippocampal metabolites/neurotransmitters were analysed longitudinally by *in vivo* magnetic resonance spectroscopy (MRS). Cognitive function was evaluated with the contextual fear conditioning test. Microglia, astrocytes, neuroblasts and interneurons were analysed by immunofluorescence.

**Results:** Approximately 50% of mice exhibited progressive post-MCAO cognitive impairment. Notably, immature hippocampal neurons in the impaired group displayed more severe aberrant phenotypes than those from the non-impaired group. Using MRS, significant bilateral changes in hippocampal metabolites such as *myo*-Inositol (*m*Ins) or N-acetylaspartic acid (NAA) were found that correlated, respectively, with numbers of glia and immature neuroblasts in the ischemic group. Importantly, some metabolites were specifically altered in the ipsilateral hippocampus suggesting its involvement in aberrant neurogenesis and remodelling processes. Specifically, MCAO animals with higher hippocampal GABA levels displayed worse cognitive outcome. Implication of GABA in this setting was supported by the amelioration of ischemia-induced memory deficits and aberrant hippocampal neurogenesis after blocking pharmacologically GABAergic neurotransmission. These data suggest that GABA exerts its detrimental effect, at least partly, by affecting morphology and integration of newborn neurons into the hippocampal circuits.

**Conclusions:** Hippocampal GABAergic neurotransmission could be considered a novel diagnostic and therapeutic target for post-stroke cognitive impairment.

## INTRODUCTION

Stroke is a major cause of death and disability worldwide, with a significant sociosanitary impact (1). In the last decades, advances in prevention and healthcare have reduced stroke patient mortality (2); therefore, stroke can now be considered a chronically disabling rather than a lethal pathology, with many survivors displaying a range of motor, cognitive and psychiatric impairments. Specifically, cognitive impairment affects at least one-third of stroke patients in the chronic phase (3), but its pathogenesis and potential treatment are unknown. Stroke drives neurogenic modifications affecting both adult neurogenic niches: the subventricular zone (SVZ) of the lateral ventricles and the subgranular zone (SGZ) of the dentate gyrus (DG) of the hippocampus (4–6). Alterations in the neurogenic process and in the integration of the newborn neurons in the latter have been proposed by us and others as causal factors contributing to cognitive impairment in different pathological scenarios including stroke (7–10). Indeed, we demonstrated that newborn neurons displayed an aberrant morphology after stroke, with opposed phenotypes at the ipsilateral and contralateral sides characterized by a different pattern of dendritic arborization probably driven by the growth of the apical dendrite. We also showed that abolition of hippocampal neurogenesis ameliorated post-stroke cognitive impairment (10). Although several underlying mechanisms could account for post-stroke aberrant neurogenesis, the specific factors that differentially modulate the ipsilateral and contralateral hippocampus leading to cognitive dysfunction are unknown. To gain insight into this process, we decided to explore post-stroke cognitive function concomitantly to longitudinal hippocampal changes in metabolites/neurotransmitters using *in vivo* non-invasive MRS together with histological assessments, and to test a pharmacological loss-of-function approach of the mechanisms identified.

## MATERIAL AND METHODS

A detailed description of the methods is provided in the Data Supplement.

### Permanent middle cerebral artery occlusion (MCAO) in mice

Mice were subjected to permanent focal cortical cerebral ischemia through the distal occlusion of the middle cerebral artery (MCA), as previously described (10).

### Magnetic resonance spectroscopy (MRS)

MRS of the hippocampus was carried out in a 7T MRI scanner (Biospec 70/30 USR, Bruker BioSpin, Ettlingen, Germany) using a transceiver mouse brain cryoProbe. Spectra were analysed with the LCModel program (11). The following metabolites were included in these analyses after checking that their %SD (estimated standard deviations expressed as percentages of the estimated concentrations relative to total creatine, i.e. Cramér-Rao lower bounds) were under 15%: γ-aminobutyric acid (GABA), glutamate (Glu), N-acetyl-aspartate (NAA), *myo-*inositol (*m*Ins), phosphorylcholine (PCh) + glycerophosphorylcholine (GPC), glutathione (GSH), and taurine.

### Treatment with the α5 subunit-selective GABA_A_ receptor (α5-GABA_A_R) inverse agonist L-655,708

L-655,708 was dissolved in dimethyl sulfoxide and frozen in aliquots. This stock was diluted 10 times in saline, to reach a final concentration of 1 mg/ml. Treatment consisted of a daily intraperitoneal injection (1mg/kg), from day 10 up to day 20 after surgery.

### Behavioural testing

Contextual fear conditioning (CFC) test was performed 7d after surgery. During conditioning, mice were placed in the chamber and, after 150s of acclimatation, they received 3-foot shocks (0.6mA, 2s duration, 1min apart). During the retrieval test, mice were placed in the chamber for 5min. Behaviour was recorded by an overhead camera and freezing (absence of movement except for breathing) quantified by an automated system (FreezeFrame4, Actimetrics).

### Image acquisition, processing and analysis

Immunofluorescence was performed on free-floating sections that were incubated overnight at 4°C with the corresponding antibodies. Acquisitions of the images were performed with a laserscanning confocal imaging system (Zeiss LSM710 at UCM; Zeiss LSM700 at CNIC). Image quantification and analysis were performed with ImageJ and Imaris.

### Statistics

Statistical analyses were performed with PRISM8.0 program (GraphPad Software Inc.) at a significance level of 95% (p<0.05).

## RESULTS

### Stroke induces progressive cognitive decline associated with hippocampal neurogenesis and bilateral remodelling

In agreement with our previous work (10), MCAO produced long-term impairment in contextual memories in mice, independently of the initial lesion extent (Fig.1A). Importantly, memory decline after stroke, observed as a reduced freezing response in the CFC test, was progressive, being undetectable at 14d after surgery but present at 35d (Fig.1B). This supports the idea that, after this type of acute ischemic cortical damage in mice, a progressive decline of the cognitive function develops, resembling that observed in stroke patients (3,12,13). Of note, not all ischemic mice displayed cognitive deficits after stroke, similar to observations in stroke patients: in another set of experiments with larger sample size, our data clearly showed two different populations of mice, where approximately 50% of them displayed a clear cognitive impairment long-term after stroke (impaired group) while the rest (non-impaired) behaved as the sham group (Fig.1C).

**Fig. 1.**
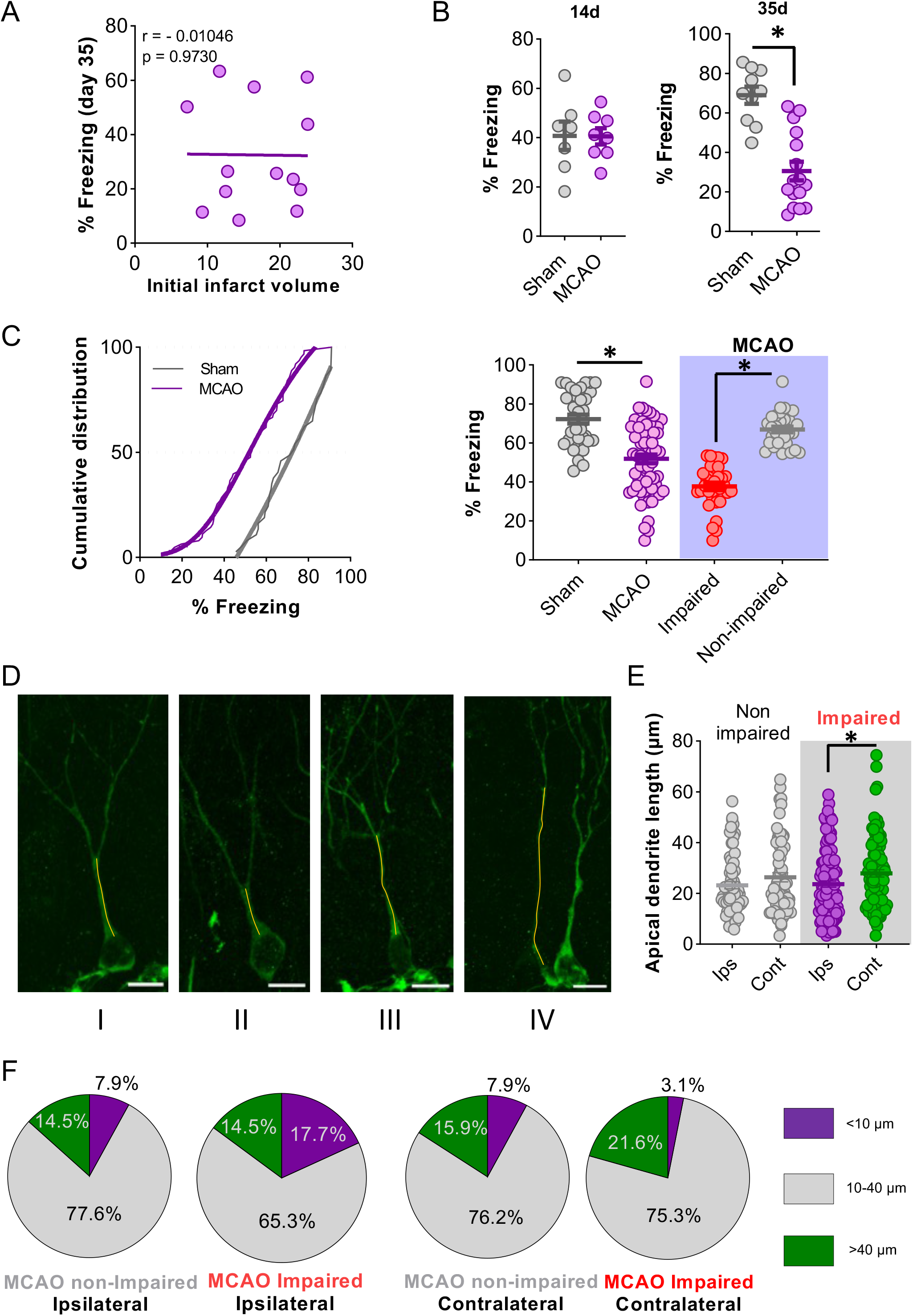
Cognitive decline after stroke is dependent of hippocampal neurogenesis and bilateral remodeling. (A) Lack of correlation between infarct volume and freezing response 35d after surgery (p>0.05). (B) Cortical stroke impairs long-term memory in mice; remote memory retention after cerebral ischemia was calculated as percentage of freezing response 14 (B left, p>0.05; sham, n=7; MCAO, n=8) and 35d (B right, *p<0.05; sham, n=10; MCAO, n=16) after surgery. (C) Cumulative distribution histogram of the freezing percentage of MCAO and sham groups 35d after surgery (left); percentage of freezing at 35d, distinguishing two subsets in the MCAO group (p<0.05; sham, n =37; MCAO, n =63) (right). (D) Representative images of a DCX^+^ cell at the ipsilateral hemisphere of a MCAO non-impaired (I) and a MCAO impaired (II), and of the contralesional hemisphere of a MCAO non-impaired (III) and a MCAO impaired (IV). (E) Comparison of apical dendrite lengths of the MCAO impaired and non-impaired on the CFC test (*p<0.05). (F) Pie charts displaying percentage of DCX^+^ cells at the ipsilesional hemisphere of the MCAO with better (n=75 DCX^+^ cells/3 mice) and worse outcome (n=121 DCX^+^ cells/4 mice), and in the contralesional hemisphere of the MCAO with better (n=63 DCX^+^ cells/3 mice) and worse outcome (n=97 DCX^+^ cells/4 mice), showing apical dendrite lengths of <10 μm, of 10-40 μm, and of >40 μm. Scale bar=10μm. Data, represented as mean±SEM, were compared by using Mann-Whitney test (B,C) or two-way ANOVA followed by Bonferroni’s test (D).

Our previous studies support that hippocampal aberrant neurogenesis is implicated in this long-term cognitive impairment (10). Consistently, 35d post-stroke, a higher percentage of newborn neurons in the ipsilateral hemisphere of the ischemic group displayed aberrant features including an ectopic location, an aberrant growth direction, an apical bipolar dendrite (Suppl. fig.1A-B) or a differential pattern of dendrite length. Furthermore, ipsilateral and contralateral newborn SGZ neurons behaved oppositely showing, respectively, either a remarkable reduction or an increase in the length of the apical dendrite (Suppl. fig.1C-D). Noteworthily, immature neurons from the impaired MCAO group displayed a more dramatic aberrant phenotype than those from the non-impaired MCAO group, strongly corroborating the involvement of hippocampal neurogenesis in cognitive decline after ischemia (Fig.1D-F).

As previously described (10), the MCAO technique used here induces infarcts restricted to the cortex, with no implication of the hippocampus (Suppl. fig.1E). In addition, no significant differences in infarct size were found between the impaired and the non-impaired groups (16.28% vs 15.46%; Suppl. fig.1E).

### Stroke induces changes in hippocampal metabolites and neurotransmitters

To gain insight into the processes underlying the differences in cognitive outcome and differential remodeling at ipsilateral and contralateral hippocampi after stroke, we performed localized *in vivo* MRS in hippocampus of sham and ischemic mice at different times after surgery (Fig. 2A). As shown (Table 1, Suppl. fig. 2), significant longitudinal changes were found *in vivo* after MCAO for different metabolites, which were seemingly independent of the initial lesion extent (Suppl. fig. 3). In particular, *m*Ins and NAA exhibited a similar pattern in both hippocampi. Regarding *m*Ins, this metabolite is a putative marker of glial cells (14), recently proposed as a prognostic marker of cognitive deficits associated with alterations in astrocyte activity in the hippocampus (15). In our hands, higher hippocampal levels of *m*Ins were associated with higher numbers and signs of activation of microglia and astrocytes, as determined by immunofluorescence at 14d for Iba-1 and GFAP and at 35d for GFAP (Fig. 2B,C,E; Suppl. fig. 4). Likewise, NAA increased bilaterally in the MCAO group (Fig. 2F). Since NAA is widely used as a neuronal marker (16), its increase 14d after surgery presumably results from the bilateral neurogenic burst that occurs after stroke (10). Accordingly, we found an increase in the number of DCX^+^ cells in both ischemic SGZ at that time (Fig. 2D,F).

**Fig. 2.**
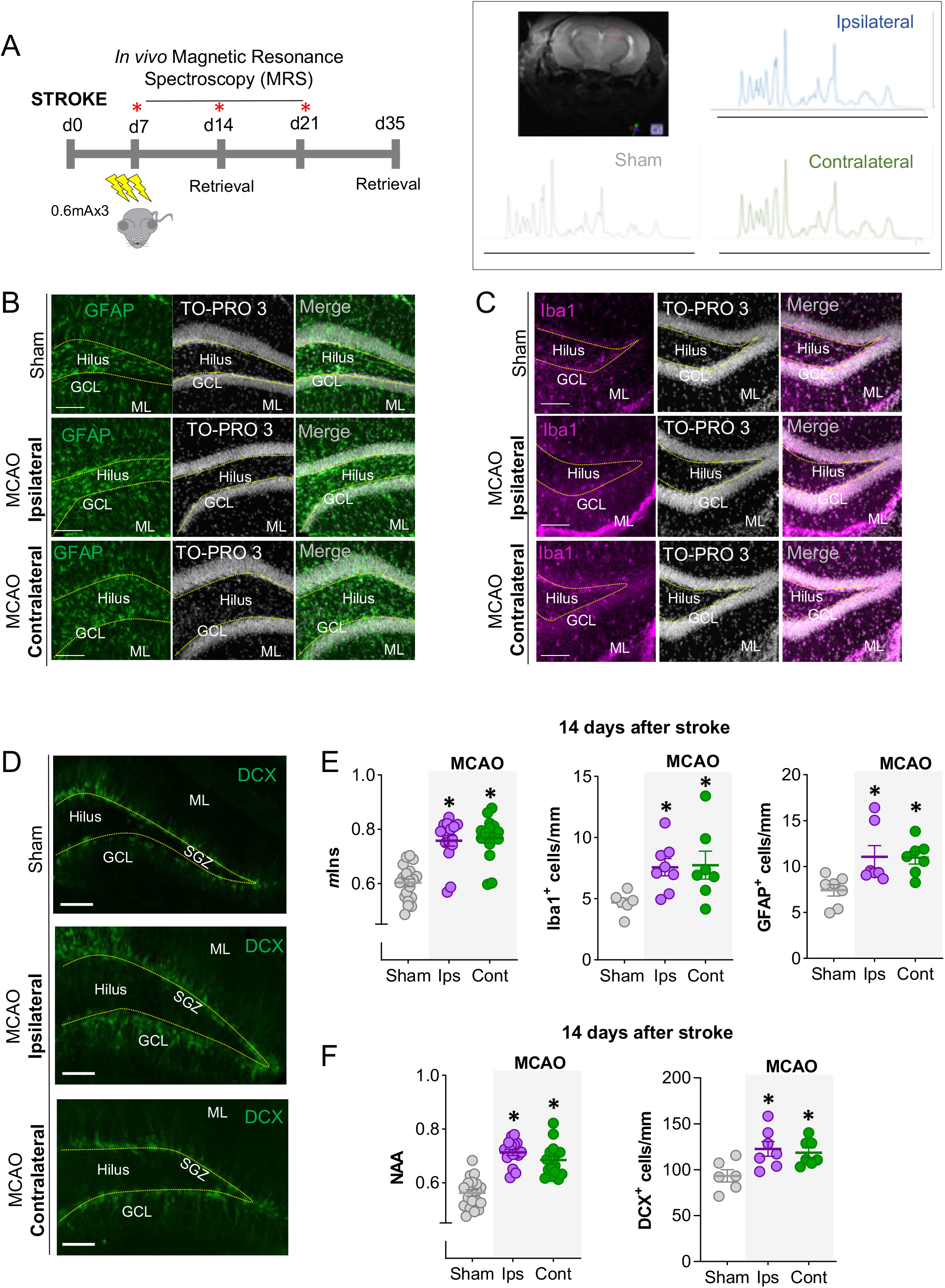
Stroke induces changes in hippocampal metabolites and neurotransmitters. (A) Experimental design of the MRS study, including CFC conditioning and retrieval times (left). Representative MRS spectra of the groups and location of acquisition voxel (right). (B-D) Representative images of GFAP^+^ (B), Iba1 ^+^ (C) and DCX^+^ cells (D) at the ipsi- and contralateral hippocampi of sham and ischemic mice 14d after surgery. TOPRO was used as a nuclear marker. (E) Hippocampal *m*Ins levels by MRS 14d after stroke (sham, n=20; MCAO, n=16; *p<0.05 vs sham; left). Density of Iba1^+^ cells and GFAP^+^ cells on the hilus and SGZ at 14d after MCAO, respectively (n=6-8; *p<0.05 vs sham) (middle and right). (F) Hippocampal NAA levels by MRS 14d after stroke (sham, n=20; MCAO, n=16; *p<0.05 vs sham) (left). Quantification of DCX^+^ cells in the DG of sham and ischemic mice 14d after the surgery (right). Scale bar=50 μm (B-C); 100μm (D). Data, represented as mean±SEM, were compared using 2-way (E-F, left) or one-way ANOVA (E middle and right; F right) followed by Tukey’s test (E-F).

**TABLE 1.** Changes in hippocampal neurometabolites after stroke. Concentration ratios (relative to creatine plus phosphocreatine) of the hippocampal neurometabolites analysed at the different time points in sham and ischemic hemispheres (* vs sham p<0.05; # vs contralesional hemisphere p<0.05; $ vs 7d; & vs 14d). Data are represented as mean±SEM. Data were compared using 2-way ANOVA followed by Tukey’s post-hoc.

| Metabolite | 7d |  |  |
| --- | --- | --- | --- |
| | Sham<br>(n=20)<br>Mean ± SEM | Ipsi<br>(n=16)<br>Mean ± SEM | Contra<br>(n=16)<br>Mean ± SEM |
| **GABA** | 0.288±0.009 | 0.296±0.009 | 0.281±0.011 |
| **Glu** | 0.881± 0.011 | 0.952±0.017* | 0.905±0.013 |
| **GPC+PCh** | 0.144±0.004 | 0.169±0.005* | 0.159±0.004* |
| **GSH** | 0.269±0.006 | 0.248±0.010 | 0.254±0.008 |
| **mIns** | 0.685±0.016 | 0.742±0.019* | 0.735±0.013 |
| **NAA** | 0.587±0.014 | 0.656±0.019* | 0.623±0.011 |
| **Taurine** | 1.098±0.015 | 1.059±0.019 | 1.040±0.012* |
| Metabolite | 14d |  |  |
| | Sham<br>(n=20)<br>Mean ± SEM | Ipsi<br>(n=16)<br>Mean ± SEM | Contra<br>(n=16)<br>Mean ± SEM |
| **GABA** | 0.240±0.012\$ | 0.303±0.007*# | 0.244±0.010 |
| **Glu** | 0.867±0.010 | 0.968±0.016*# | 0.910±0.015 |
| **GPC+PCh** | 0.138±0.004 | 0.159±0.004* | 0.148±0.002* |
| **GSH** | 0.251±0.005 | 0.252±0.008 | 0.270±0.008 |
| **mIns** | 0.603±0.014\$ | 0.758±0.020* | 0.768±0.019* |
| **NAA** | 0.562±0.012 | 0.713±0.011*\$ | 0.685±0.015*\$ |
| **Taurine** | 1.097±0.021 | 1.044±0.018 | 1.050±0.013 |
| Metabolite | 21d |  |  |
| | Sham<br>(n=20)<br>Mean ± SEM | Ipsi<br>(n=16)<br>Mean ± SEM | Contra<br>(n=16)<br>Mean ± SEM |
| **GABA** | 0.274±0.020 | 0.320±0.009*# | 0.251±0.018 |
| **Glu** | 0.887±0.018 | 0.980±0.018*# | 0.912±0.019 |
| **GPC+PCh** | 0.147±0.006 | 0.170±0.006*# | 0.148±0.005 |
| **GSH** | 0.272±0.007 | 0.242±0.007* | 0.260±0.007 |
| **mIns** | 0.660±0.017& | 0.772±0.021* | 0.771±0.017* |
| **NAA** | 0.594±0.021 | 0.740±0.028*\$ | 0.676±0.018* |
| **Taurine** | 1.113±0.012 | 1.101±0.022 | 1.088±0.020 |

Furthermore, other metabolites were altered in the ipsilateral vs. the contralateral side such as glutamate and GABA, or vs. the sham group, such as GPC+PCh (Table 1; Suppl. fig.2). Among these, the high levels of GABA and glutamate in the ipsilesional vs. the contralesional hemisphere and the sham group remained elevated at 14 and 21d after stroke (Fig. 3A, Table 1, Suppl. fig.2). However, only GABA levels at 14d after stroke showed a significant negative correlation with freezing, in such a way that MCAO animals with higher hippocampal GABA levels displayed worse cognitive outcome 35d after the injury (Fig. 3B; Suppl. figs. 5-7). These data strongly support that GABA levels are involved in the development of post-stroke cognitive impairment.

**Fig. 3.**
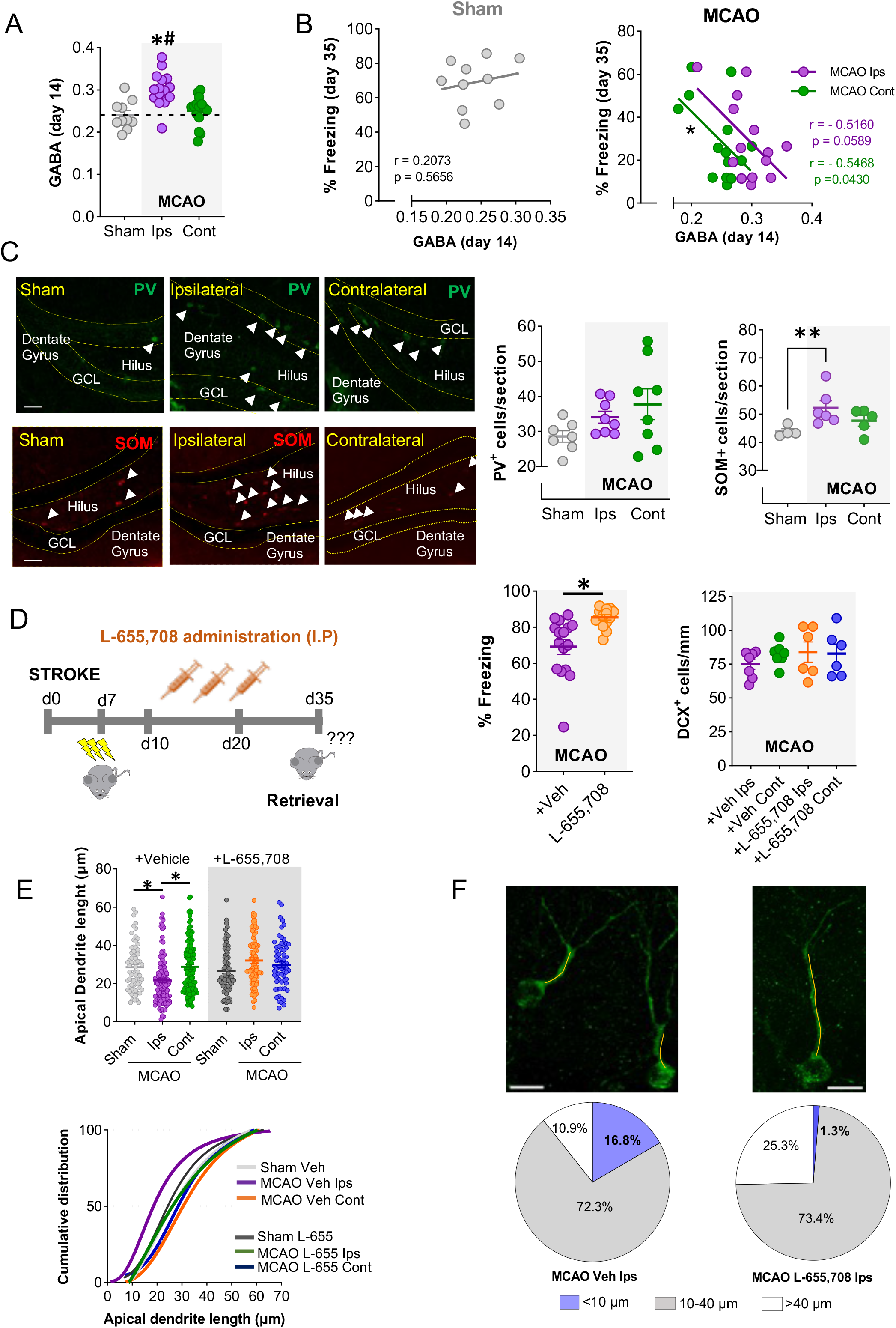
Post-stroke memory impairment is reduced by blocking GABA_A_ receptors. (A) Hippocampal GABA levels by MRS 14d after stroke (*p<0.05 vs sham, #p<0.05 vs contralesional; sham, n=20; MCAO, n=16). (B) Comparison of Pearson’s correlations between GABA levels 14d after surgery and percentage of freezing 35d after surgery (sham, n=10, Pearson’s r=0.2073, p=0.5656; ipsilesional MCAO, n=14, Pearson’s r=0.5160, p=0.0589; contralesional MCAO, n=15, Pearson’s r=0.5468, p<0.05). (C) Representative images of parvalbumin (PV)^+^ (green) and somatostatin (SOM)^+^ interneurons (red) (left). Immunoreactivity of PV^+^ interneurons per section (SGZ, hilus, GL, ML, CA1, CA2, CA3) analysed 14d after surgery (n=7-8, p>0.05; middle). Immunoreactivity of SOM^+^ interneurons per section (hilus and SO) analysed 14d after surgery (*p<0.05 vs sham) (right). (D) Experimental design for antagonising GABAergic activation using the α5-GABAAR inverse agonist L-655,708 during the hippocampal post-stroke neurogenic peak (left). Percentage of freezing response in vehicle- and L-655,708-treated sham and MCAO mice 35d after surgery (*p<0.05; sham vehicle, n=6; L-655,708 sham, n=5; vehicle MCAO, n=15; L-655,708 MCAO, n=12; middle). Quantification of the number of DCX^+^ cells/mm on the groups treated with vehicle or L-655,708, 35d after surgery (right). (E-F) Morphological features of newborn neurons in vehicle- and L-655,708-treated sham and MCAO mice, 35d after surgery. In (E), mean apical dendrite length (top) and histogram (bottom) showing the frequency distribution of the apical dendrite length of groups treated with vehicle (p<0.05 MCAO ipsilesional vs sham, and MCAO ipsilesional vs MCAO contralesional) or L-655,708 (p>0.05). (F) Top: representative images of DCX^+^ cells comparing the apical dendrite length of ipsilesional neuroblasts from vehicle- or L-655,708-treated MCAO mice. Bottom: pie charts display the percentage of DCX^+^ cells at the ipsilesional hemisphere of MCAO vehicle (n=119 DCX^+^ cells/5mice) or MCAO L-655,708 (n=79 DCX^+^ cells /5mice) showing apical dendrite lengths of <10 μm, of 10-40 μm, and of >40 μm. Scale bar=50μm (C); 10μm (F). Data, represented as mean±SEM, were compared using 2-way (A,D-F) or one-way ANOVA (C) followed by a Tukey’s (A,C,D right,E,F) or Bonferroni’s test (D middle). Correlation analysis was assessed by Pearson’s (B).

### Stroke differentially affects the immunoreactivity of hilar GABAergic interneurons at the dentate gyrus

We next quantified by immunofluorescence the most representative and prevalent populations of GABAergic neurons in the hippocampus, i.e., parvalbumin- and somatostatin-expressing interneurons. While no significant changes were found in the number of immunoreactive parvalbumin^+^ interneurons neither at 14 nor at 35d (Fig. 3C and Suppl. fig. 8, respectively), we observed a significant increase of ipsilesional hilar immunoreactive somatostatin^+^ neurons vs. sham (Fig. 3C), probably indicating an increased expression of somatostatin in those interneurons after stroke.

### Post-stroke memory impairment is reduced by blocking GABA_A_ receptors

As increased GABA levels and cognitive impairment after ischemic injury might be related, we hypothesized that blocking GABAergic neurotransmission might ameliorate ischemia-induced hippocampal deficits, at least in part, by modulating the neurogenic response. Given the predominant expression of GABA_A_ receptors in hippocampus, specifically of the α5-GABA_A_R (17,18) and the literature supporting their role in cognitive function (19), we decided to use the selective α5-GABA_A_R inverse agonist L-655,708 (20–23), administered from post-surgery day 10 to 20 coinciding with the post-stroke neurogenesis peak (10), to explore whether interference with the elevated GABAergic neurotransmission could reverse post-stroke long-term cognitive decline. Confirming our hypothesis, L-655,708 treatment abolished cognitive decline after MCAO (Fig. 3D, left). The drug did not have any effect on non-conditioned mice (Suppl. fig. 9A-B) or sham mice (Suppl. fig 9C, left), ruling out stroke-independent mechanisms for the drug and supporting that up-regulated GABA levels underlie post-stroke cognitive impairment.

Finally, we wondered whether the beneficial outcome on cognitive performance induced by L-655,708 could be associated with effects on the neurogenic process and/or the morphology of the newborn neurons. To check this, we analysed both neuroblasts number and morphology by DCX staining. We did not observe major differences in the numbers of DCX^+^ neuroblasts between vehicle- and L-655,708-treated groups (Fig. 3D and Suppl. fig. 9C, right), thus discarding effects at the proliferation stage. However, L-655,708 treatment significantly reduced the number of newborn immature neurons displaying an aberrant morphology. Notably, the α5-GABA_A_R inhibitor decreased ipsilateral aberrant phenotype of newborn neurons, as demonstrated by a reduction in the percentage of immature neurons displaying a shortening of the apical dendrite length (Fig. 3E-F and Suppl. fig. 9D). Overall, our data very plausibly support that the elevated ipsilateral hippocampal GABA levels observed after stroke can contribute to cognitive hippocampal deficits, at least in part, by affecting the morphology and integration of the newborn neurons into the hippocampal circuits.

## DISCUSSION

Clinical studies show that at least a third of the survivors of cerebrovascular events suffer post-stroke cognitive decline (3), thus reflecting its magnitude and clinical importance. However, mechanisms involved remain obscure, largely due to the complex and heterogeneous pathophysiology of stroke subtypes. Here, we have used a well characterised model of distal stroke in mice, which presents a very reproducible cortical lesion and hippocampal neurogenesis-dependent cognitive alterations (10), to gain insight into the mechanisms underlying post-stroke cognitive impairment. Our results reveal that hippocampus-dependent memory deficits and underlying aberrant SGZ neurogenesis are associated, at least partly, with GABA elevations that we have found, using *in vivo* localized MRS, in the ipsilesional hippocampus of mice after distal cortical stroke, suggesting that hippocampal GABAergic overactivation could be a diagnostic and therapeutic target for post-stroke cognitive impairment.

Stroke has been shown to significantly stimulate neurogenesis in the adult DG, including the MCAO models by either permanent or transient MCAO (24–27). Therefore, remote cortical infarcts not affecting the hippocampal formation are able to promote a significant augmentation in hippocampal neurogenesis. Although there is still some controversy regarding the functional impact of increased neurogenesis in the DG after cerebral ischemia, recent studies have shown that hippocampal newborn neurons display morphological alterations that might hinder their proper integration into the hippocampal circuits (7,8,10,28,29) and, therefore, underlie hippocampus-dependent memory deficits. Importantly, in agreement with observations in humans (3), we have found that approximately one half of the ischemic mice presented long-term memory deficits after stroke (impaired group), while the rest did not (non-impaired memory group). This finding provided a convenient setting to explore the mechanisms accounting for such differences. Indeed, our data demonstrate that only the impaired group displayed significant differences in dendritic morphology when comparing ipsi- vs contralesional hippocampi, strongly supporting the implication of aberrant neurogenesis in post-stroke memory deficits.

The late onset (35d after the surgery) of the cognitive deficit after MCAO is similar to the progressive deterioration observed in stroke patients (30), and consistent with the time required for the development of an aberrant neurogenesis and the subsequent maladaptive hippocampal remodelling. However, the mechanisms responsible for this process are unknown. Therefore, we decided to carry out a longitudinal characterisation of hippocampal neurotransmitters/metabolites after stroke, by means of non-invasive *in vivo* MRS, a technique useful to study *in vivo* ischemic stroke and dementia in animal models and clinical settings (31–33). Interestingly, MRS results showed two types of changes: some bilateral, i.e. occurring at both hippocampi, and some unilateral, i.e. mainly associated with the ipsilesional side.

As for the bilateral changes, those involving *m*Ins and NAA were the most evident ones. *Myo*-Inositol is a metabolite primarily synthesized in glial cells. In fact, its bilateral increase herein reported at 14 and 21d post-MCAO was in agreement with glial cell numbers found at 14 and 35d. Traditionally, glial activation after stroke has been associated with the acute phase, during the first 48-72 hours, in areas close to the infarct core, and as part of the inflammatory response (34). However, we have found high levels of both glial activation and *m*Ins at later time points and far from the injured area. As regards microglia, its increase in number could result from phagocytosis of the excess in neural stem cells (35), or from the synaptic pruning of newborn neurons (36). On the other hand, a bilateral increase in astrocytes contributing to increased *m*Ins values could arise from the asymmetric division of neuroblasts towards astrocytic cells (37). NAA, used as a neuronal marker (16), also showed a significant bilateral increase 14d after the injury, in agreement with an increase in the number of neuroblasts in both ischemic SGZ at that time. Further studies are necessary to confirm these associations.

More importantly, considering the association between unilateral differences with post-stroke cognitive dysfunction at the CFC, we focused our attention on ipsi- vs contralesional differences. Several metabolites showed this pattern, in particular GABA (14 and 21d post MCAO), glutamate (14 and 21d post MCAO), and GPC+PCh (21d post MCAO). Notably, higher hippocampal GABA levels 14d after MCAO correlated with worse memory deficits. Since this significant correlation was not detected for any other of the molecules studied, we felt prompted by the appealing hypothesis of the involvement of GABA in hippocampus-dependent memory deficits after ischemic injury. This possibility was strengthened by the previously described importance of GABA in the neurogenic mechanism (38–41). Interestingly, GABA, the classic inhibitory neurotransmitter in mature neurons, is excitatory in immature neurons, neuroblasts and neural stem and progenitor cells (42). Whereas the excitatory GABA signalling during early neurogenesis is important for activity/experience-induced regulation of neural stem cells quiescence, neural progenitor cells proliferation, neuroblast migration and newborn neuronal maturation/functional integration, inhibitory GABA signalling allows for the sparse and static functional networking essential for learning/memory development and maintenance (40,42,43), a duality necessary for a correct neural network development. Consequently, an altered GABAergic influence such as the tonic increased transmission observed in our setting is likely to affect negatively the above mentioned processes, impairing long-term potentiation (22) and contributing to the learning and memory deficits observed.

Tonic inhibition is mainly mediated by GABA_A_ receptors containing α5 subunit especially in hippocampus (44, 45). Thus, we wondered whether pharmacological interference with the GABAergic over-activation using L-655,708, a benzodiazepine inverse agonist selective for α5-GABA_A_R (46), could reverse memory deficits in our model. Indeed, we verified satisfactorily that L-655,708 treatment improved memory retrieval, as shown by an increased freezing response in the treated mice 35d days after MCAO. This finding is consistent with previous ones in which both genetic knock-down or pharmacological blockade of α5-GABAAR ameliorated cognitive behaviour, including hippocampus-dependent performance (20–22,47).

To further confirm our hypothesis, we studied if the beneficial effect of the α5-GABAAR inverse agonist on cognitive performance was associated with a reversal of the aberrant features of the neurogenic process. Indeed, the treatment induced a recovery of the morphological phenotype, with ipsilateral apical dendrites lengths not significantly different either from the contralesional or the sham ones. All these data together strongly support the notion that an over-activated GABAergic neurotransmission underlies the aberrant newborn neuronal morphology and the subsequent hippocampus-dependent memory deficits after stroke in mice. Supporting our data, it was recently shown that an increased astrocytic GABA release in hippocampus interfered with neurotransmission resulting in poor memory performance in a mouse model of Alzheimer disease (48). In this context, we cannot discard that, in addition to neurons, astrocytes could play a role in the modulation of GABA levels in post-stroke hippocampus (49, 50).

We also characterised the two major groups of GABAergic interneurons, specifically parvalbumin^+^ and somatostatin^+^ interneurons. At 14d after MCAO, time at which GABA levels correlate with hippocampal memory deficits, of parvalbumin^+^ immunoreactive interneurons did not show any differences between groups. Notably, immunoreactivity of the somatostatin^+^ subset showed a significant increase in the ipsilesional hemisphere. The enhanced immunoreactivity at this stage, between the genesis of the newborn neurons and their integration in the hippocampal circuits, might indicate an increased activation of these interneurons as a result of their modulating function. Likewise, an increase of somatostatin^+^ interneurons in the DG was observed in a model of temporal lobe epilepsy (51). Somatostatin^+^ interneurons have been described to control the formation of neuronal assemblies during memory acquisition, modulate synaptic plasticity and contribute to synchronization of neuronal ensembles and the generation of gamma rhythms (51–53). Importantly, increasing their activity during training has been shown to lower the freezing response (54) and they were also found to be the main contributors to enhanced GABA synaptic activity in genetic mouse models of Huntington’s disease (55).

Overall, we hereby show that the development of hippocampus-dependent memory deficits after cortical stroke depend on the establishment of a morphologically aberrant neurogenesis associated with a tonic elevation of ipsilesional GABA at the hippocampus. The inhibition of both memory deficits and maladaptive neurogenesis by an inverse agonist of α5-containing GABA_A_ receptors suggests that hippocampal GABAergic overactivation is a novel therapeutic target for post-stroke cognitive impairment, offering a novel predictor and therapeutic approach for the chronic phase of stroke patients.

## Author contributions

CTL, AGC, MIC, JDP, JMGS, IL and MAM designed the research studies. CTL, AGC, JDP, SVR, TJF, JMGS, EFV and MB conducted the experiments and/or acquired the data. CTL, MIC, AGC, JDP, JMGS, OH, MSB and MAM contributed to the analysis and/or interpretation of the results. CTL, MIC, AGC and MAM wrote the manuscript, which all authors reviewed and approved.

## Acknowledgements

This work was supported by grants from Spanish Ministry of Science and Innovation (MCIN) PID2019-106581RB-I00 (MAM), from Leducq Foundation for Cardiovascular Research TNE-19CVD01 (MAM, MSB) and TNE-21CVD04 (MAM, IL), and from Instituto de Salud Carlos III (ISCIII) and co-financed by the European Development Regional Fund “A Way to Achieve Europe” PI20/00535 and RICORS-ICTUS RD21/0006/0001 (IL). CNIC is supported by ISCIII, MCIN and ProCNIC Foundation, and is a Severo Ochoa Center of Excellence (CEX2020-001041-S). The microscopy experiments were performed in Unidad de Microscopía e Imagen Dinámica, CNIC, ICTS-ReDib, co-funded by MCIN/AEI /10.13039/501100011033 and FEDER “Una manera de hacer Europa” (#ICTS-2018-04-CNIC-16). Part of the research work included in this publication has been carried out in the ReDIB ICTS infrastructure BioImaC, MCIN.

