## Supplemental Material for "IPSILESIONAL HIPPOCAMPAL GABA IS ELEVATED AND CORRELATES WITH COGNITIVE IMPAIRMENT AND MALADAPTIVE NEUROGENESIS AFTER CORTICAL STROKE IN MICE"

### SUPPLEMENTAL METHODS

#### Animals

All experimental protocols were in accordance with the guidelines of the Animal Welfare Committee of UCM, CNIC and Consejería de Medio Ambiente y Ordenación del Territorio de la Comunidad de Madrid (RD 53/2013; PROEX 047/16) following European directives 86/609/CEE and 2010/63/EU, and reported in accordance with ARRIVE guidelines (Animal Research: Reporting of In Vivo Experiments) (1). The article adheres to the Transparency and Openness Promotion Guidelines. Study data are available upon request. Experimental groups until April 2021 were housed at the animal facility of UCM and from May 2021 at the animal facility of CNIC. Animals were kept under standard conditions of temperature and humidity and a 12h-light-dark cycle with food and water available *ad libitum*.

#### Permanent middle cerebral artery occlusion (MCAO) in mice

Mice were anesthetized with isoflurane 1.5%-2% (UCM) or sevoflurane 2-3% (CNIC) in a mixture of 80% air/20% oxygen; body temperature was maintained at physiological levels with a heating pad during the process. The MCA was occluded by ligature of the trunk just before its bifurcation between the frontal and parietal branches with a 9-0 suture, in combination with the occlusion of the ipsilesional common carotid artery, as described (2). Mice in which the MCA was exposed but not occluded served as sham-operated controls.

#### Magnetic resonance spectroscopy (MRS)

MRI (magnetic resonance imaging) was acquired before MRS in the same session for each animal. MRS of the hippocampus was carried out at 7, 14 and 21d after surgery in a Biospec7T USR70/30 (Bruker, Ettlingen) using a Transceiver Mouse Brain CryoProbe (300 MHz, Bruker BioSpin MRI, Ettlingen). To ensure the best animal welfare, MRI was performed at low resolution (256×256 pixels, 1 pixel=0.0586mm). Briefly, MRI acquisition protocol included an initial flash sequence (TR:100ms, TE:2.5ms, FOV:3cm, matrix:128×128 pixels) to obtain three orthogonal slices. For homogeneity, we acquired a field map sequence in all brain. Then, anatomical images were taken in three positions to localize the voxel for the spectrum. The sequence used was T2-turborare (TR:2500ms, TE:33ms, FOV:1.5cm, matrix:256×256 pixels, Z=400μm). 1H-PRESS sequence was used to acquire the spectrum (TR:2000ms, TE:11ms, number of averages:512, FOV:3.80mm3; time of experiment:17min). Spectra were analysed with the LCModel program (11) using a simulated metabolic profile based on TE = 11ms PRESS sequence at 7T. No filtering was applied to the FIDs. The LCModel analysis was performed on each spectrum from the chemical shift range of 0.0-4.0ppm. The concentration ratios relative to creatine plus phosphocreatine were used for the analysis. The following metabolites were included: γ-aminobutyric acid (GABA), glutamate (Glu), N-acetyl-aspartate (NAA), inositol (*m*Ins), phosphorylcholine (PCh) + glycerophosphorylcholine (GPC), glutathione (GSH) and taurine.

Animals were anesthetised with isoflurane (3% for induction and 1-1.5% for maintenance); respiration and body temperature were continuously monitored (SA Instruments, NY).

**Infarct volume measurement**

Infarct size was determined by MRI 24 to 48h after MCAO using a Biospec BMT 47/40 (Bruker; UCM) or a 7T MRI (Agilent-Varian; CNIC). Infarct volume was calculated using ImageJ software from the T2-weighted images as previously described (2).

**Contextual fear conditioning (CFC) test**

Since enduring CFC memories has been ascribed to the hippocampus, we used this test to evaluate hippocampal function. During the course of this study, two different conditioning chambers were used: for experiments shown in Table 1, Figs. 1,2,3A-C and Suppl. figs. 1-8, CFC was performed in test chambers (AM1000 Avoidance Station, Kinder Scientific) of 31×24×21cm and shock-grid floors with bars 3.2mm in diameter spaced 7.9mm apart. For the experiments shown in Fig. 3D-F and Suppl. fig. 9, CFC was performed in a conditioning chamber (Actimetrics, 80014AT) of 18.7x20.06x20.1cm and shock-grid floors with bars 4mm in diameter spaced 8.15mm apart. To ensure maximum reduction in external light and sound, the conditioning chamber was located inside a cubicle (Actimetrics, 83020AT) of 45.75x45.75x55.1cm.

In both cases, CFC was performed 7d after surgery. During conditioning, mice were placed in the chamber and, after 150s of acclimatation, they received 3-foot shocks (0.6mA, 2s duration, 1min apart). Intensity levels in both apparatuses were validated with an amperemeter. One minute after the last shock, mice were returned to their housing cages. During the retrieval test, mice were placed in the chamber for 5min. Behaviour was recorded by an overhead camera and freezing (absence of movement except for breathing) quantified by an automated system (FreezeFrame 4, Actimetrics).

**Histology**

*Tissue preparation*

Mice were perfused transcardially with phosphate buffer (0.1M) followed by 4% paraformaldehyde (PFA). Brains were post-fixed in PFA and transferred to 30% sucrose. Coronal sections (30µm) were cut using a microtome (Leica SM2000R) and stored in cryoprotective solution. Series of 1:10 coronal sections spaced 300µm apart were used.

*Immunofluorescence*

Immunofluorescence was performed on free-floating sections that were incubated overnight at 4°C with the following antibodies: rabbit anti-DCX (1:250, Abcam, ab18723); chicken anti-GFAP (1:500, Thermo Scientific, PA1-10004); rabbit anti-Iba1 (1:250, Wako, 011-27991); mouse anti-somatostatin (1:500, GeneTex, GTX71935). For parvalbumin $(PV)^+$ interneurons staining, incubation with mouse anti-PV (1:750, Sigma-Aldrich, P3088) lasted 48h. The secondary antibodies used were goat anti-rabbit IgG (H+L) biotinylated (1:500, Vector Laboratories, BA-1000); streptavidin Alexa Fluor 488 conjugate (1:500, Thermo Scientific, S32354); goat Alexa

Fluor anti-chicken 488 (1:500, Thermo Scientific, A11039); donkey Cy3 anti-rabbit (1:500, Millipore, AP182C); goat Alexa Fluor 488 anti-mouse (1:500, Thermo Scientific, A10680); goat Alexa Fluor 555 anti-mouse (1:500, Thermo Scientific, A32727).

*Image acquisition, processing and analysis*

Acquisitions were performed with a laser-scanning confocal imaging system (Zeiss LSM710 at UCM; Zeiss LSM700 at CNIC). Image quantification and analysis were performed with ImageJ and Imaris. Doublecortin (DCX)$^+$ cells, Iba1$^+$ cells, GFAP$^+$ cells, PV$^+$ and somatostatin (SOM)$^+$ interneurons were counted in confocal Z-stack images for which 3-5 serial sections (30µm) per animal spaced 300µm apart were analysed. The number of DCX$^+$ cells was quantified in the SGZ. The number of Iba1$^+$ cells and GFAP$^+$ cells was quantified in the hilus and the SGZ of the DG. The number of PV$^+$ was quantified in the different regions of the hippocampus (SGZ, hilus, molecular and granular layer, CA1, CA2 and CA3). The number of SOM$^+$ cells was quantified in the hilus and in the *stratum oriens* (SO). Counting of the positive cells was carried out manually. Data of Iba1$^+$ cells, GFAP$^+$ cells and DCX$^+$ cells are expressed in number of cells/mm, and the data of the interneurons are expressed as number of cells/section.

For the morphological study of the astrocytes, ten GFAP$^+$ cells were randomly selected from the SGZ in each section and their morphological complexity was analysed using the "Sholl analysis" plugin of ImageJ. For the morphological study of the neuroblasts at the SGZ, DCX$^+$ cells were analysed using the *Filaments* and *Sholl analysis* plugins from Imaris. Specifically, the analysis of the total dendritic length, the apical dendrite length and the number of intersections was carried out. In the analysis of the aberrant DCX$^+$ cells, the number of bipolar cells, of cells located ectopically in the hilus or in the upper part of the granular layer, and of cells growing in abnormal directions were counted. Researchers blinded to the experimental conditions performed confocal imaging and data quantification.

**Statistics**

Statistical analyses were performed using the PRISM8.0 program (GraphPad Software Inc.) at a significance level of 95% ($p<0.05$). All data are presented as group mean values ± standard error of the mean (SEM). Normality tests were performed. Pairwise comparisons were evaluated using the Student's t-test or the Wilcoxon-Mann-Whitney non-parametric test. Multiple comparisons were evaluated using one or two-way ANOVA, followed by Bonferroni's, Tukey's or Dunn's post-hoc. For the correlation analysis, Pearson's test was performed.

**SUPPLEMENTARY FIGURES**

**Suppl. fig. 1. Characterization of post-stroke aberrant neurogenesis.** (A) Schematic representation of the aberrant neuroblasts types: (I) normal morphology; (II) bipolar neuroblast, without apical dendrite and with two or more filaments from the soma; (III) neuroblast with altered growth direction; (IV) ectopic neuroblast in the upper part of the granular layer; (V) ectopic neuroblast in the hilus. (B) Aberrant $DCX^+$ cells quantification (left); quantification of the bipolar type (*p<0.05) (right). (C) Pie charts display percentage of $DCX^+$ cells in each group showing apical dendrite lengths of <10μm, of 10-40 μm, and of >40μm (sham, n=229 $DCX^+$ cells/8 mice; MCAO ipsilesional n=169 $DCX^+$ cells/6 mice; MCAO contralesional, n=142 $DCX^+$ cells/6 mice). (D) Representative images of apical dendrites of the mentioned lengths range. (E) Representative MRI images of the infarcted hemisphere 24h after the surgery of a non-impaired MCAO and an impaired MCAO (top); percentage of infarct volume of the non-impaired MCAO group and the impaired MCAO group (bottom).

**Suppl. fig. 2. Concentration ratios (relative to creatine plus phosphocreatine) of the hippocampal neurometabolites analysed longitudinally by MRS at 7, 14 and 21d after MCAO** (*p<0.05 vs sham; #p<0.05 vs contralateral).

**Suppl. fig. 3**. **Lack of correlation between infarct volume and *in vivo* levels of the different neurometabolites analysed** (p>0.05). Correlation analysis was assessed by Pearson's. Metabolite levels are expressed as concentration ratios relative to creatine (plus phosphocreatine).

**Suppl. fig. 4**. **Analysis of hippocampal glial cells**. (A) Density of microglia ($Iba1^+$ cells; left) and astrocytes ($GFAP^+$ cells; right) on the hilus and SGZ 35d after the surgery (n=6-8; *vs sham, p<0.05). (B-C) Sholl analysis of the $GFAP^+$ cells in the SGZ of sham and MCAO mice. B: Representative image of $GFAP^+$ cells 14d after MCAO (left). A significant interaction between distance from the soma and the number of intersections was observed ($F_{(40,\ 2709)}$=2.285, p<0.0001; *p<0.05 sham vs. ipsilesional MCAO and #p<0.05 sham vs contralesional MCAO; sham, n=40 cells/7 mice; ipsilesional MCAO, n=42 cells/7 mice; contralesional MCAO, n=50 cells/8 mice) (middle). Total length of the $GFAP^+$ cells in the SGZ of sham and MCAO mice (right). C: Representative image of $GFAP^+$ cells 35d after the surgery (left). No significant interaction between distance from the soma and the number of intersections was observed ($F_{(38,\ 2300)}$=1.210, p>0.05; #p<0.05 sham vs MCAO contralesional; sham, n=43 cells/5 mice; ipsilesional MCAO, n=36 cells/6 mice; contralesional MCAO, n=39 cells/6 mice) (middle). Total length of $GFAP^+$ cells in the SGZ of sham and MCAO mice (right). Scale bar=10 μm. Data are represented as mean±SEM. Data were compared using one-way ANOVA followed by Tukey's post-hoc (A) or using Kruskal-Wallis followed by Dunn's (B, C, right) or 2-way ANOVA followed by Tukey's (B, C, middle).

**Suppl. fig. 5 Hippocampal metabolites determined by MRS 7d after surgery**. No correlation was found between the levels of the neurometabolites analysed in sham and MCAO mice 7d after surgery and the percentage of freezing response 28d after the conditioning (sham, n=10; ipsilesional MCAO, n=14-15; contralesional MCAO, n=14-15; p>0.05). Correlation analysis was assessed by Pearson's.

**Suppl. fig. 6 Hippocampal metabolites determined by MRS 14d after surgery**. Except from GABA, no correlation was found between the levels of the neurometabolites analysed in sham and MCAO 14d after surgery and the percentage of freezing response 28d after the conditioning (sham, n=10; ipsilesional MCAO, n=14-15; contralesional MCAO, n=14-15; p>0.05). Correlation analysis was assessed by Pearson's.

**Suppl. fig. 7 Hippocampal metabolites determined by MRS 21d after surgery**. No correlation was found between the levels of the neurometabolites analysed in sham and MCAO 14d after surgery and the percentage of freezing response 28d after the conditioning (sham, n=10; ipsilesional MCAO, n=14-15; contralesional MCAO, n=14-15; p>0.05). Correlation analysis was assessed by Pearson's.

**Suppl. fig. 8 Analysis of hippocampal interneurons.** Immunoreactivity of PV+ interneurons per section (SGZ, hilus, GL, ML, CA1, CA2, CA3) analysed 35d after surgery (p>0.05) (left). Immunoreactivity of the SOM+ interneurons per section (hilus and SO) analysed 35d after surgery (p>0.05) (right). Data, represented as mean±SEM, were compared using one-way ANOVA followed by Tukey's post-hoc.

**Suppl. fig. 9. Post-stroke memory impairment is reduced by blocking $GABA_A$ receptors: supplemental data.** (A) Experimental design. (B) Percentage of freezing response in L-655,708-treated naive compared with the vehicle- and L-655,708-treated shams (sham vehicle, n=6; sham L-655,708, n=5; non-conditioned naive, n=5; *p<0.05). (C) Percentage of freezing response in vehicle- and L-655,708-treated sham mice 35d after surgery (p>0.05; sham vehicle, n=6; L-655,708 sham, n=5) (left). (D) Pie charts display percentage of $DCX^+$ cells in the control SGZ and the contralesional SGZ, treated with vehicle or L-655,708, showing apical dendrite lengths of <10 μm, of 10–40 μm, and of >40 μm (sham vehicle, n=68 $DCX^+$ cells /3 mice; sham L-655,708 n=71 $DCX^+$ cells/3 mice; MCAO contralesional vehicle, n=131 $DCX^+$ cells/5 mice; MCAO contralesional L-655,708, n=65 $DCX^+$ cells /4 mice). Data were compared using one-way ANOVA (B) or 2-way ANOVA (C) followed by Tukey's post-hoc.

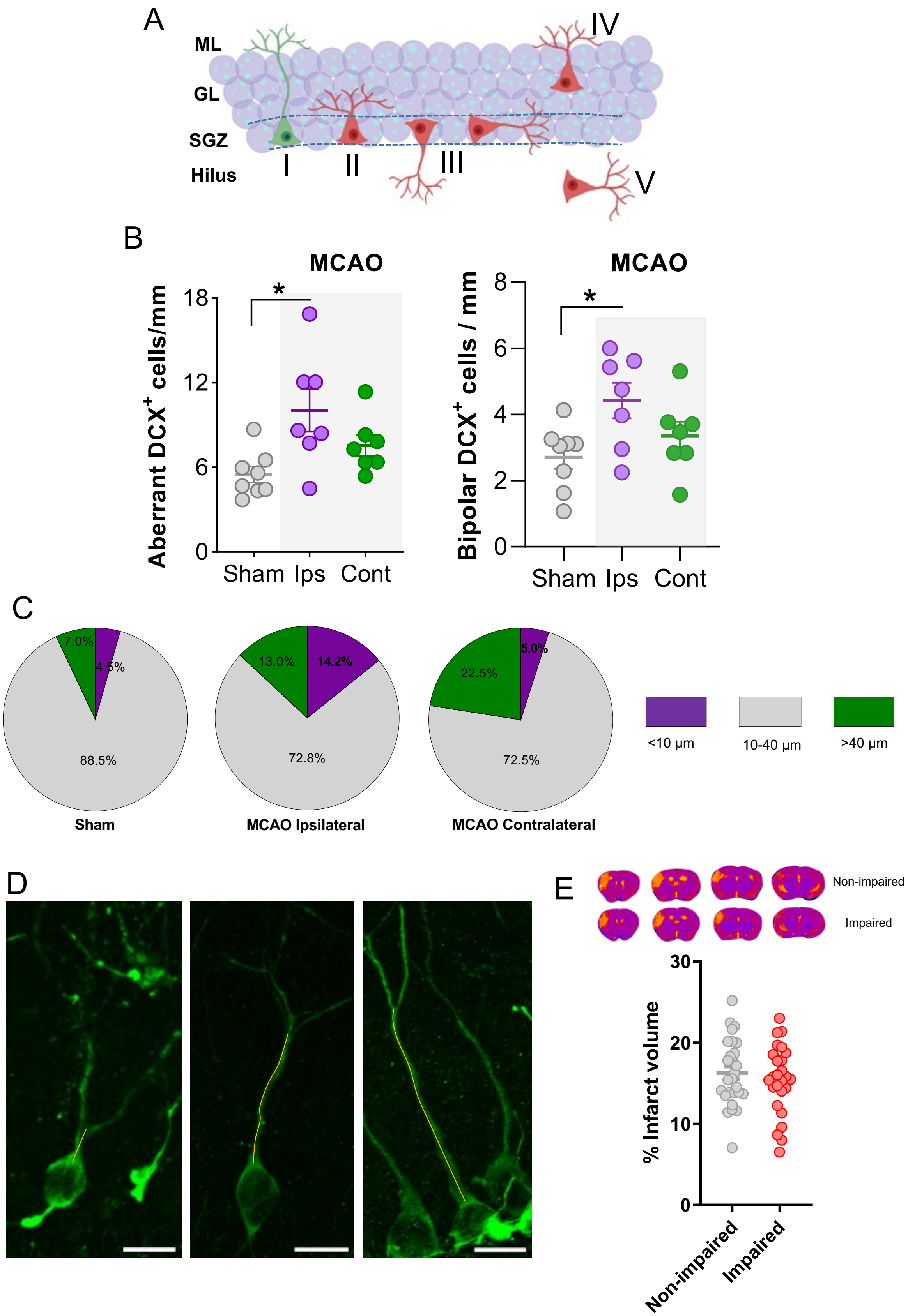


**Supplemental fig. 1**

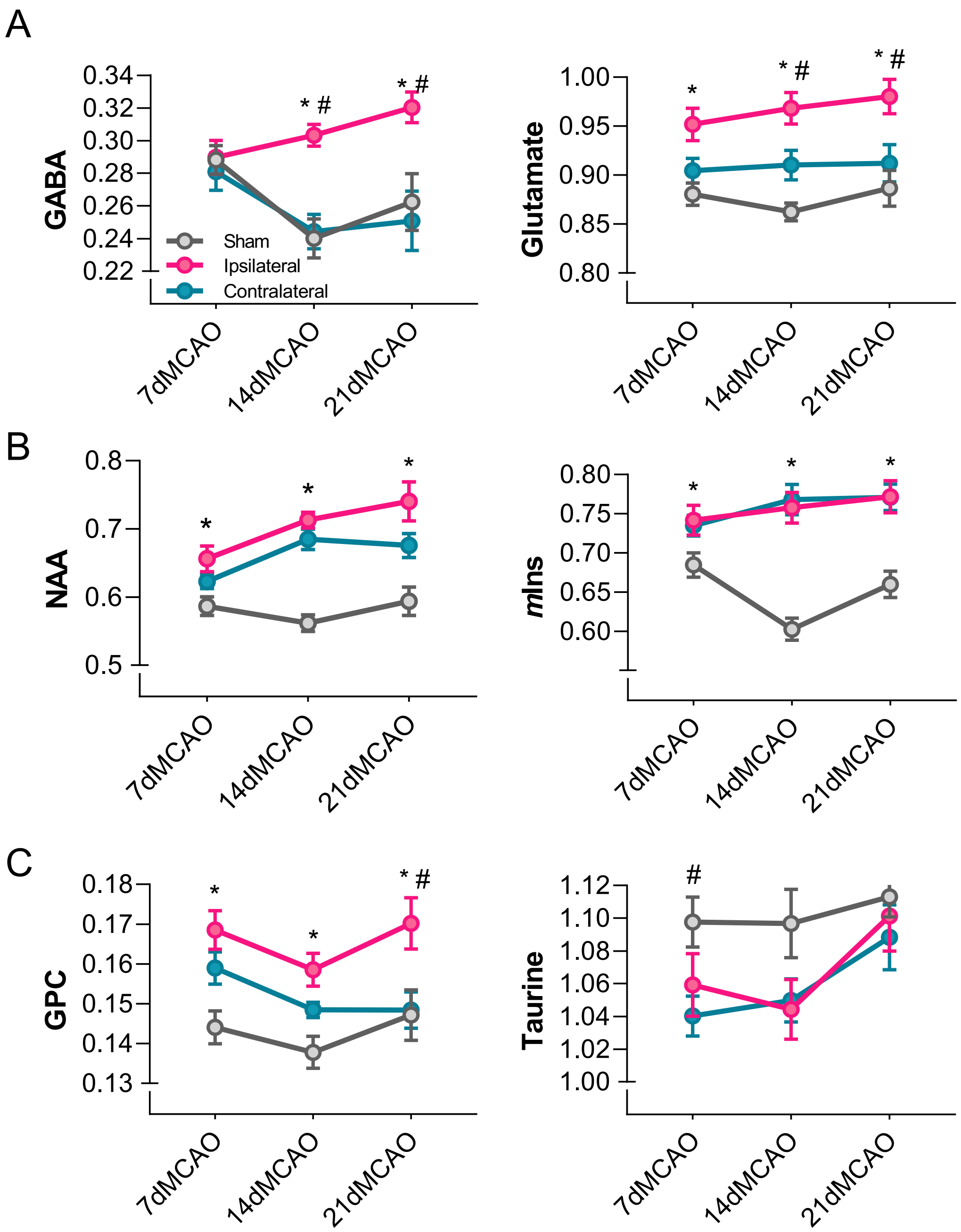


**Supplemental fig. 2**

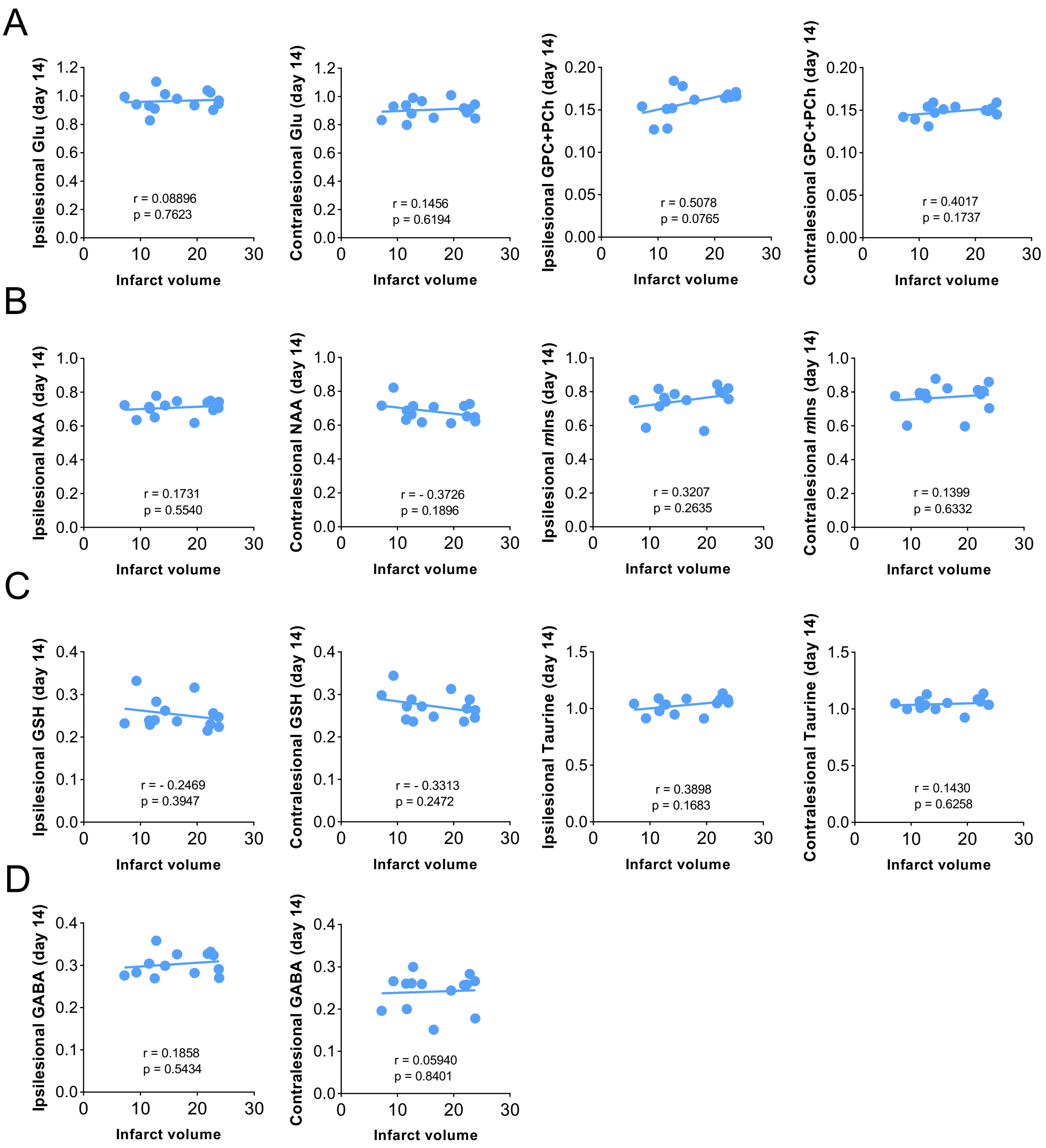


**Supplemental fig. 3**

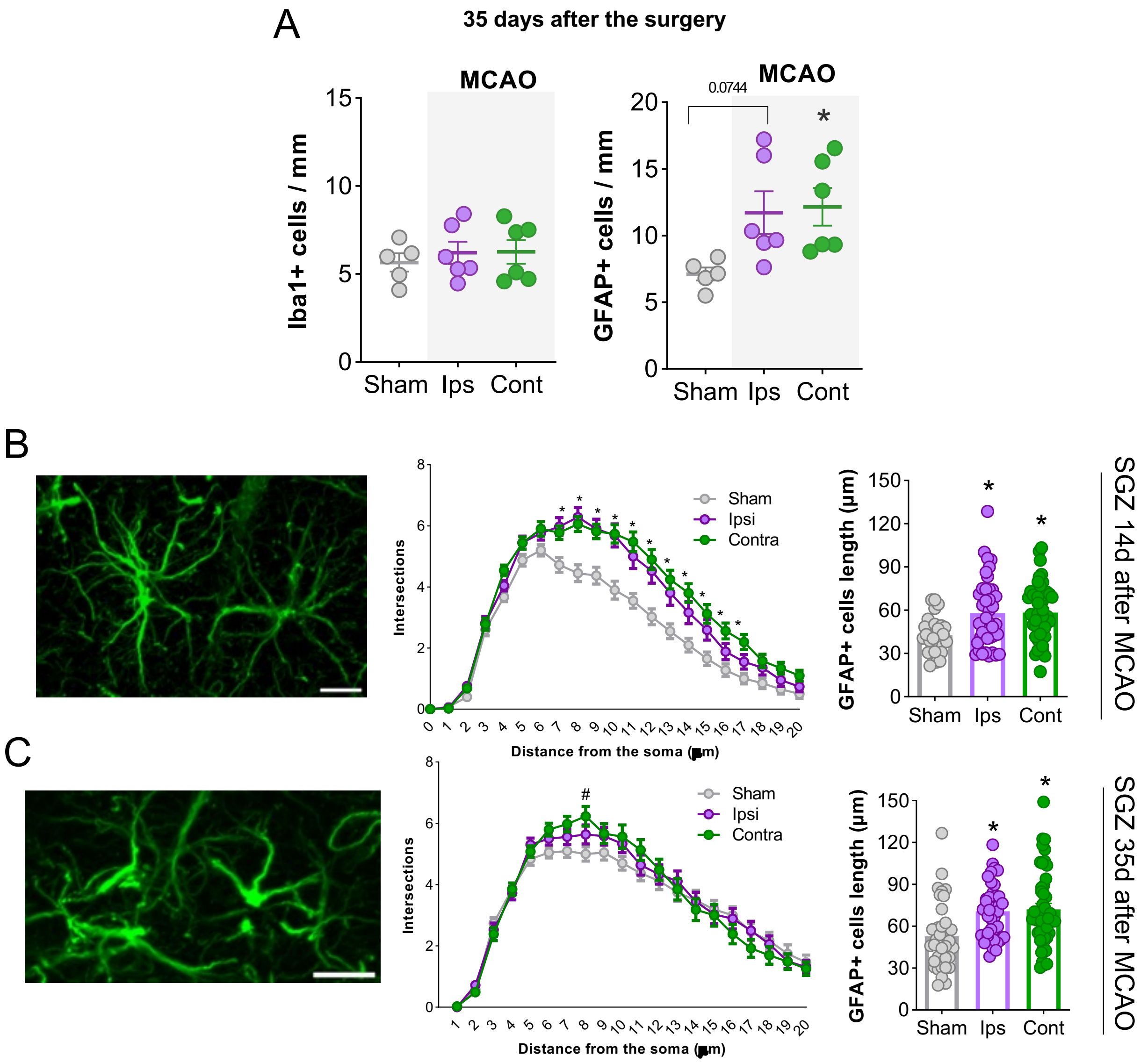


**Supplemental fig. 4**

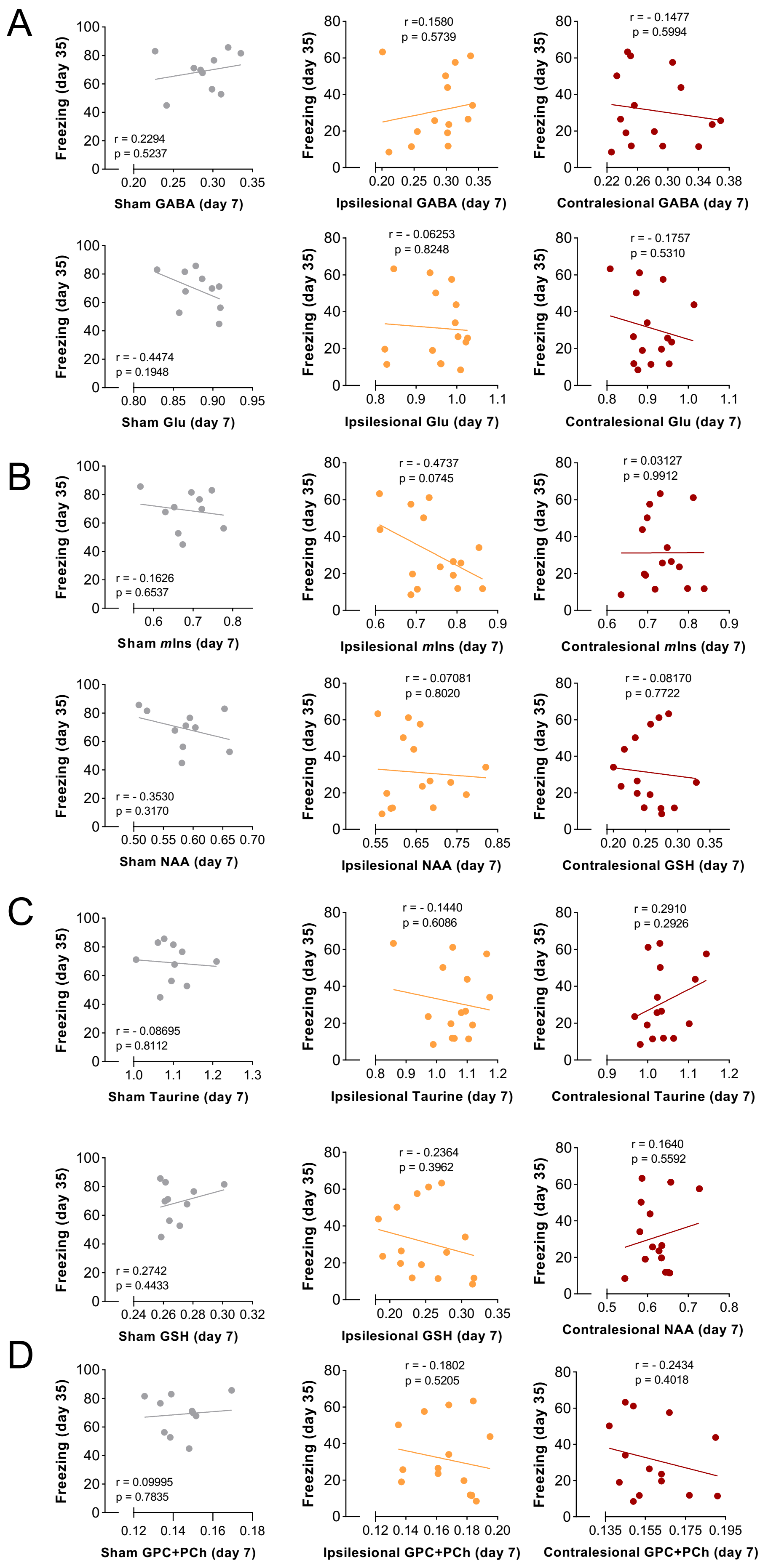


**Supplemental fig. 5**

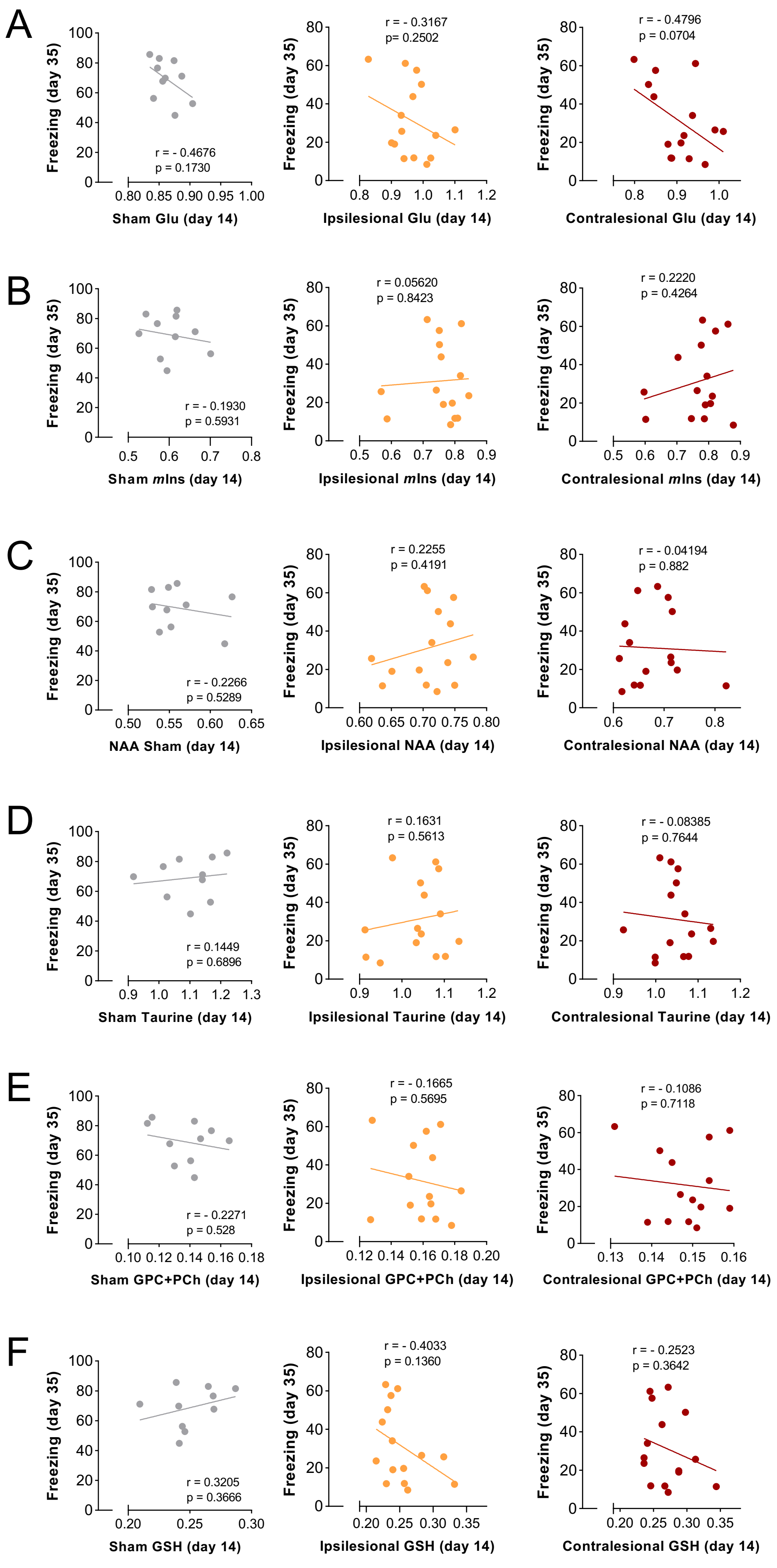


**Supplemental fig. 6**

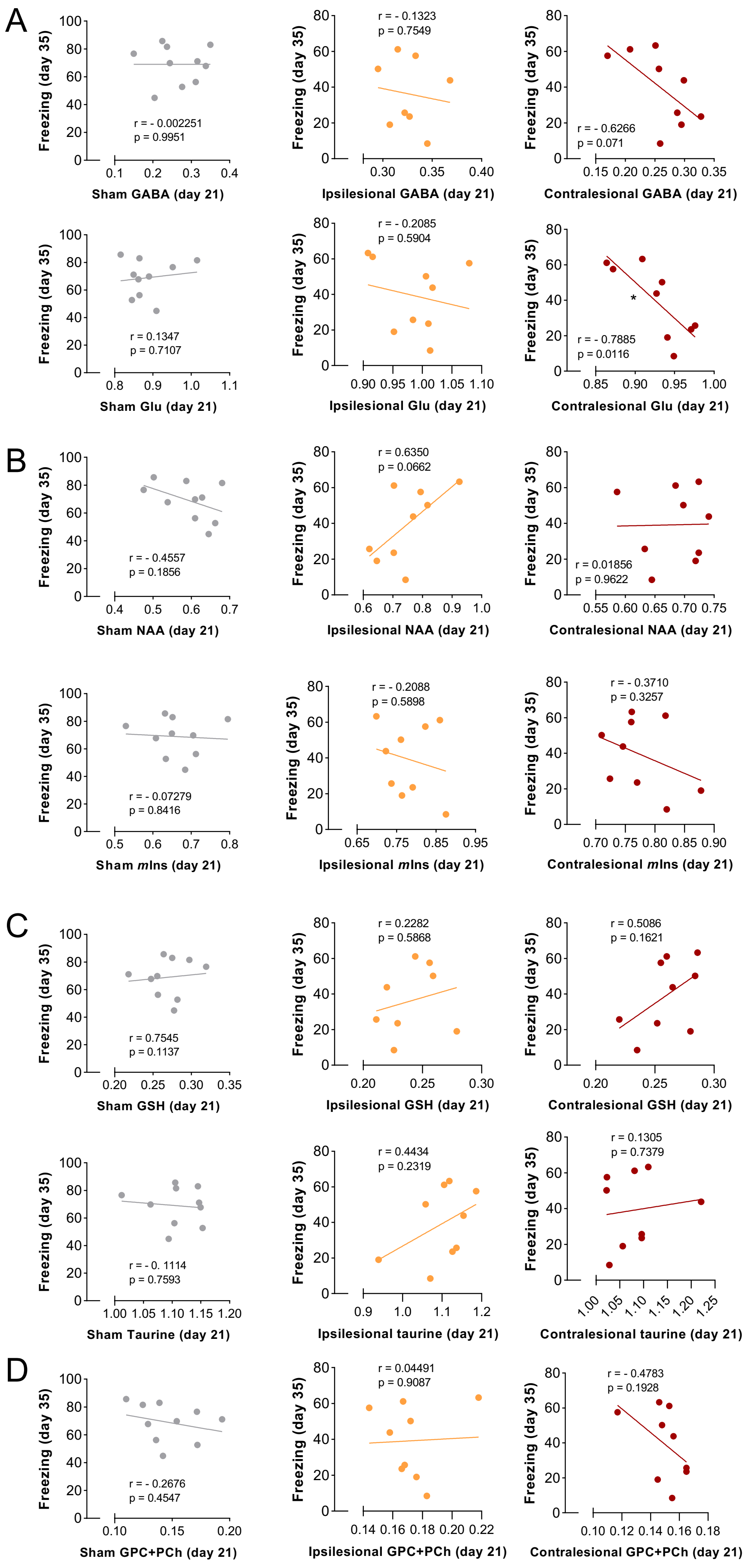


**Supplemental fig. 7**

**35d**

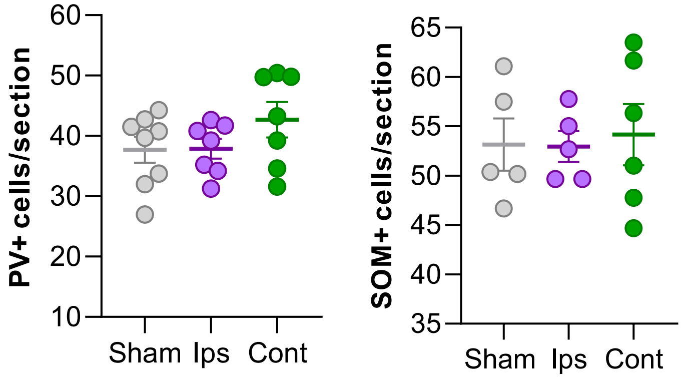


**Supplemental fig. 8**

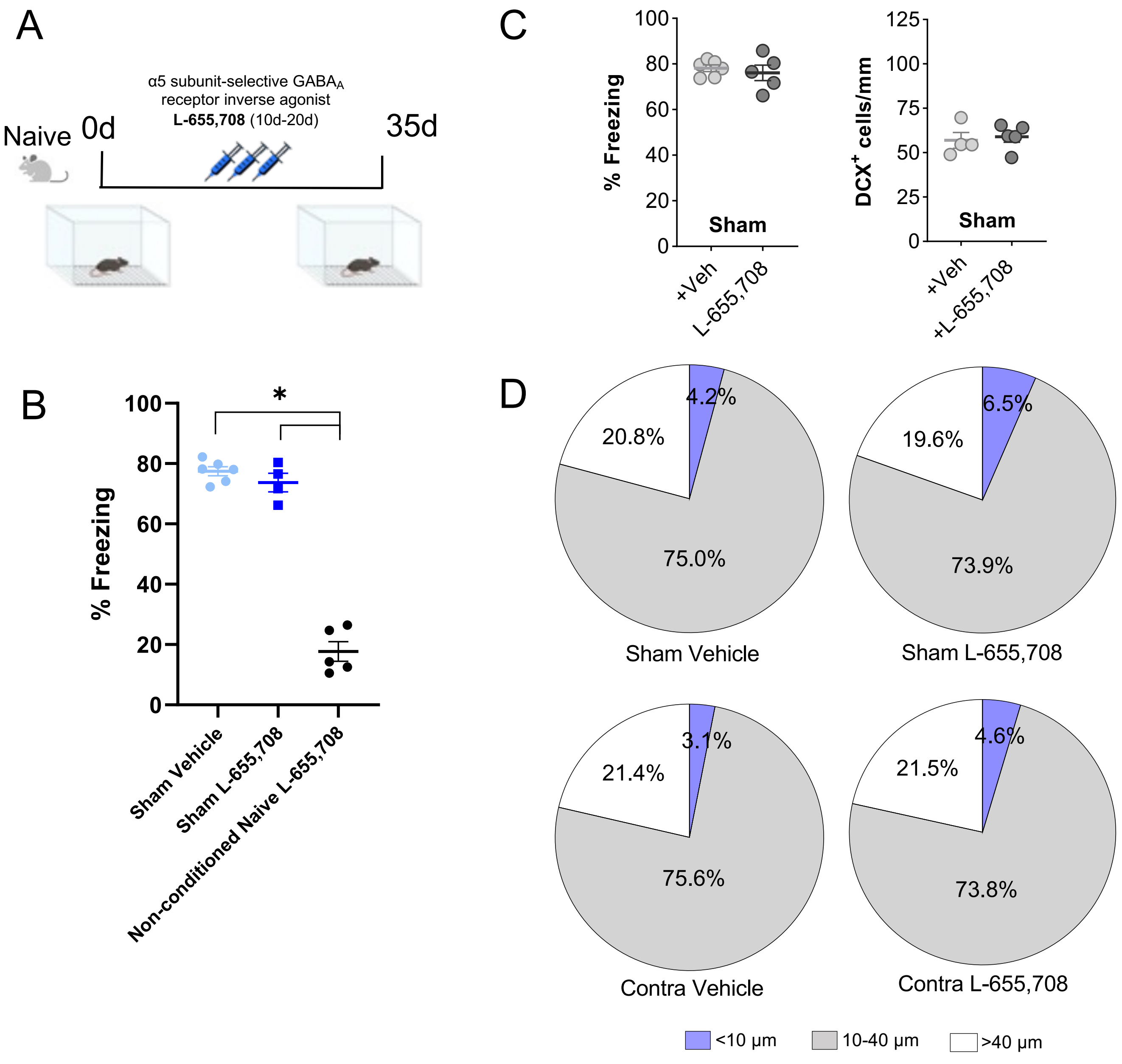


**Supplemental fig. 9**
